# Folding landscape of a parallel G-quadruplex

**DOI:** 10.1101/526780

**Authors:** Robert D. Gray, John O. Trent, Sengodagounder Arumugam, Jonathan B. Chaires

## Abstract

Circular dichroism and stopped-flow UV spectroscopies were used to investigate the thermodynamic stability and the folding pathway of d[TGAG3TG3TAG3TG3TA2] at 25 °C in solutions containing 25 mM KCl. Under these conditions the oligonucleotide adopts a thermally stable, all-parallel G-quadruplex topography containing three stacked quartets. K^+^-induced folding shows three resolved relaxation times, each with distinctive spectral changes. Folding is complete within 200 s. These data indicate a folding pathway that involves at least two populated intermediates, one of which seems to be an antiparallel structure that rearranges to the final all-parallel conformation. Molecular dynamics reveals a stereochemically plausible folding pathway that does not involve complete unfolding of the intermediate. The rate of unfolding was determined using complementary DNA to trap transiently unfolded states to form a stable duplex. As assessed by 1D-^1^H NMR and fluorescence spectroscopy, unfolding is extremely slow with only one observable rate-limiting relaxation time.

**TOC GRAPHICS:** 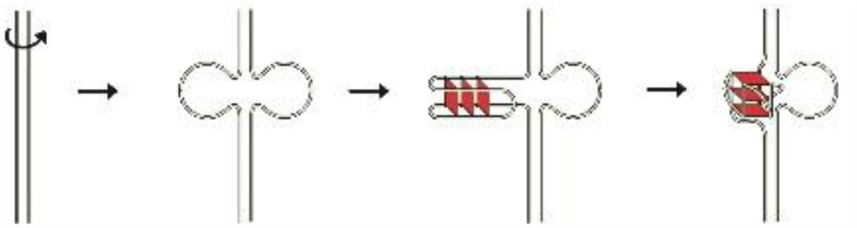

---

The folding mechanism of G-quadruplex (G4) forming DNA has been the subject of numerous studies over the past decade. Most of the kinetic studies carried out either in solution^1–5^ or by various single-molecule methods^6–17^ have focused on oligonucleotide models of the human telomere G4 consisting of variations of the sequence d[(GGGTTA)_n_] where n ≥ 4^18–19^. Depending on the exact sequence and identity of added monovalent cation, such telomeric oligonucleotides generally fold into either antiparallel “basket” or “chair” forms or into “hybrid” 3+1 antiparallel G4 topologies in multi-step reactions with spectroscopically distinct intermediates. For example, K^+^-induced folding of the human telomere sequence, d[A(G_3_TTA)_3_G_3_] initially forms an ensemble of hairpins which collapse to an antiparallel structure within ~2 s at room temperature. Subsequently, the antiparallel structure slowly rearranges to a mixture of hybrid-1 and hybrid-2 topologies over a period of hours.^1, 4–5^ Unfolding of telomeric oligonucleotides in the presence of excess complementary DNA is multiphasic and relatively slow (time constant ~2000 s) at room temperature.^5, 20–21^ However, to date there are limited solution studies on the mechanism of folding of G4 DNA folded in an all-parallel topology. The purpose of the current investigation was to apply spectroscopic methods to determine the folding and unfolding pathways for a parallel G4 structure.

To accomplish this objective, we chose the G4-forming oligonucleotide d[TGAG_3_TG_3_TAG_3_TG_3_TA_2_] (that we call “1XAV” after the Protein Data Bank entry), a well-characterized truncated sequence variant of the *c-MYC* promoter DNA. In the presence of K^+^, NMR studies show that this and related sequences fold into a G4 structure of exceptional thermal stability in which the four G_3_-tracts are in a parallel configuration.^3, 22–23^. Recently a high-resolution (2.35Å) crystal structure of the same sequence folded into a parallel G4 structure was reported ^24^. A single-molecule study using magnetic tweezers was used to estimate an unfolding rate constant of ~10^−6^ s^−1^ for a related *c-MYC* sequence folded into a parallel G4 structure, with the assumption that folding was a two-state mechanism with no intermediates.^25^ Hysteretic forward and reverse thermal denaturation transitions were used to estimate the folding and unfolding rate constants as a function of temperature for a several *c-MYC* sequence variants, again assuming a simple two-state mechanism.^3^ For some variants that showed hysteresis (but not 1XAV), unfolding rates at physiological temperature of ~200-400 s^−1^ were reported in 2 mM KCl. No kinetic intermediates were considered. We report here thermodynamic and kinetic studies that describe the folding of 1XAV in more detail, revealing populated intermediates within its folding landscape.

Figures 1, S1 and S3 show the results of equilibrium titration and thermal denaturation studies of 1XAV. Upon addition of KCl, the initially unfolded oligonucleotide adopts a structure with a CD spectrum characterized by a maximum at 265 nm, a change in sign at 253.5 nm, and a minimum at 243 nm, consistent with formation of an all-parallel quadruplex topology.^26–28^ The titration curve shown in Figure 1A, derived from more complete spectral data in Figure S1, shows that folding is not a simple two-state process. A phenomenological fit of the titration curve to a biphasic dose-response curve yielded estimates of 67 [61.5;72.8] and 73 [51.2;104.3] μM for the KCl concentrations at half-maximal saturation of each phase, where the brackets indicate the 95% confidence interval. The corresponding Hill coefficients for the two transitions are 2.1 [1.5;2.8] and 0.6 [0.3;0.8]. The very low amplitude of the second transition contributes to the uncertainty of its parameter estimated, although inclusion of the second transition is required for an optimal fit to the experimental data as judged by the randomness of residuals for the fit.

**Figure 1.**
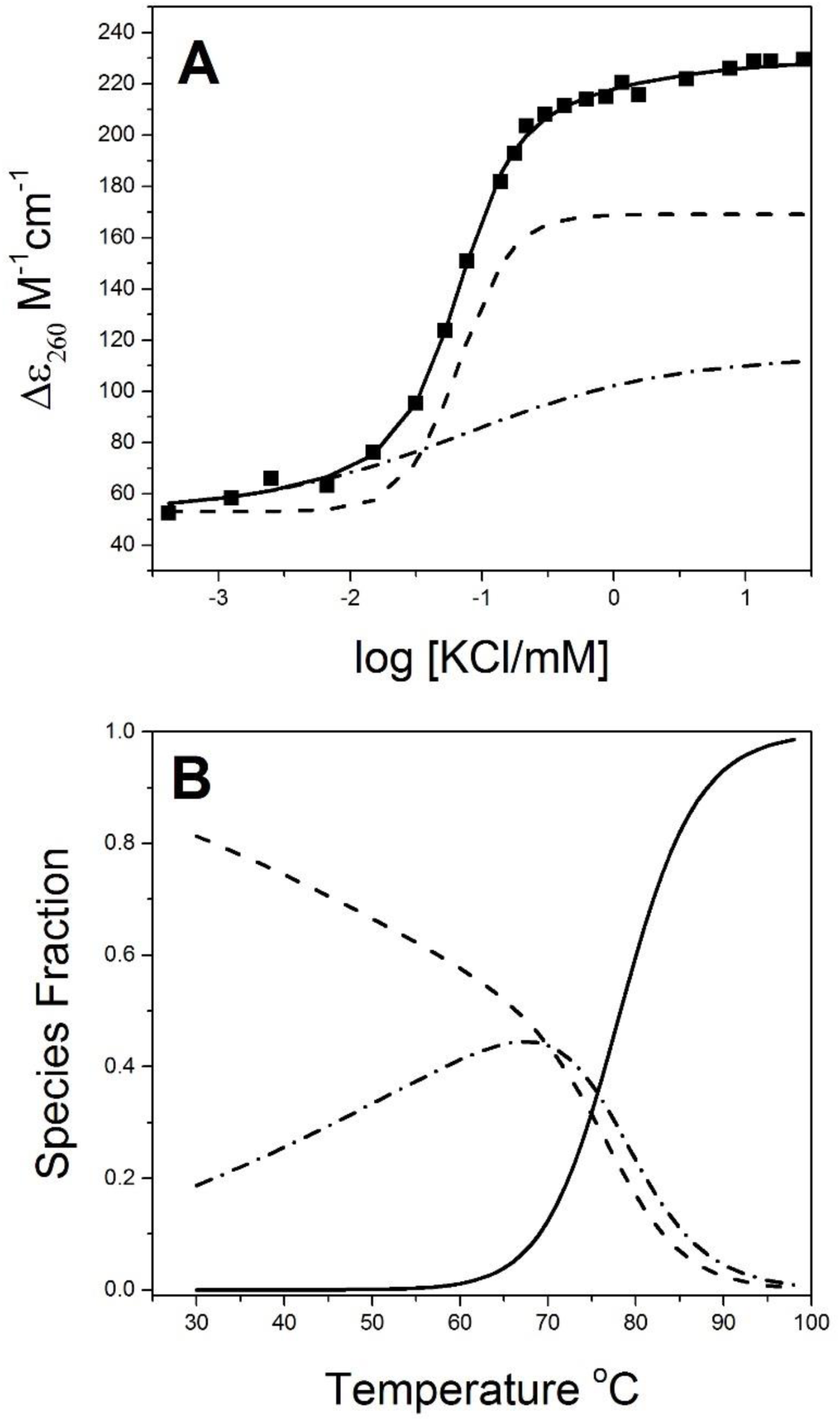
Equilibrium studies of 1XAV folding. (A) KCl titration of 1XAV. The data were fit to a double Hill equation (solid line) to obtain transition midpoints and Hill coefficients. (B) Species distribution plot derived from SVD analysis of thermal denaturation experiments. The dashed line shows the disappearance of the initial folded form, the dot-dash line shows the transient formation of an intermediate and the solid line shows the formation of the unfolded strand.

Singular value decomposition was used to analyze thermal denaturation experiments ^29^ as shown in Figure S2 and summarized in Figure 1B. The results show that thermal unfolding is not a simple two-state process, and that a populated intermediate must be included in the reaction mechanism. The two steps in the unfolding process in 25 mM KCl derived by fitting the temperature dependent family of CD spectra as previously described ^29–30^are characterized by transition melting temperatures of 57.6 ± 6.9 and 78.3 ± 2.1 °C, with van’t Hoff enthalpy changes of 19.6 ± 8.7 and 53.9 ± 3.4 kcal mol^−1^, respectively. From these thermodynamic values, the overall free energy of folding of 1XAV at 20 °C is estimated to be -11.2 ± 1.3 kcal mol^−1^ (assuming no heat capacity change). The errors are standard deviations of the mean for six independent replicate melting experiments. The complete thermodynamic profile for 1XAV folding is shown in Figure S2. Folding is driven by a favorable enthalpy and is opposed by an unfavorable entropic contribution.

The results of single wavelength stopped-flow CD experiments for data collected at 295 nm are summarized in Figure 2. To capture the complete kinetic profile, kinetic traces using the same solutions were collected in separate experiments for 2 s (Figure 2A) and 60 s (Figure 2B). The 2-s data sets could by accurately fit to a single exponential from which a relaxation time of approximately 0.3 s could be determined. The amplitude of this step depended strongly on the wavelength monitored (Figure S3). As shown in Figure 2B, the 60 s data were clearly biphasic, showing the rapid increase in ellipticity during the initial 1-2 s, followed by a slower decrease in ellipticity over the ensuing 60 s with a relaxation time of approximately 9 s. Figure 2C shows a kinetic difference spectrum for the 2-s relaxation that was generated by measuring the kinetics of CD changes at different wavelengths (see Figure S3 for individual experiments). This kinetic difference CD spectrum shows that the folding intermediate(s) initially formed exhibit a spectrum with a maximum near 295 nm that is reminiscent of an antiparallel G4 structure, perhaps mixed with some parallel or hybrid structure. These data show unequivocally that folding is not a simple two-state process, and instead proceeds through at least one intermediate form with distinctive spectral properties. (It is interesting to note that since the duration of a single eye blink in 0.1-0.4 s ^31–32^, the initial folding event is literally over in a blink of the eye.)

**Figure 2.**
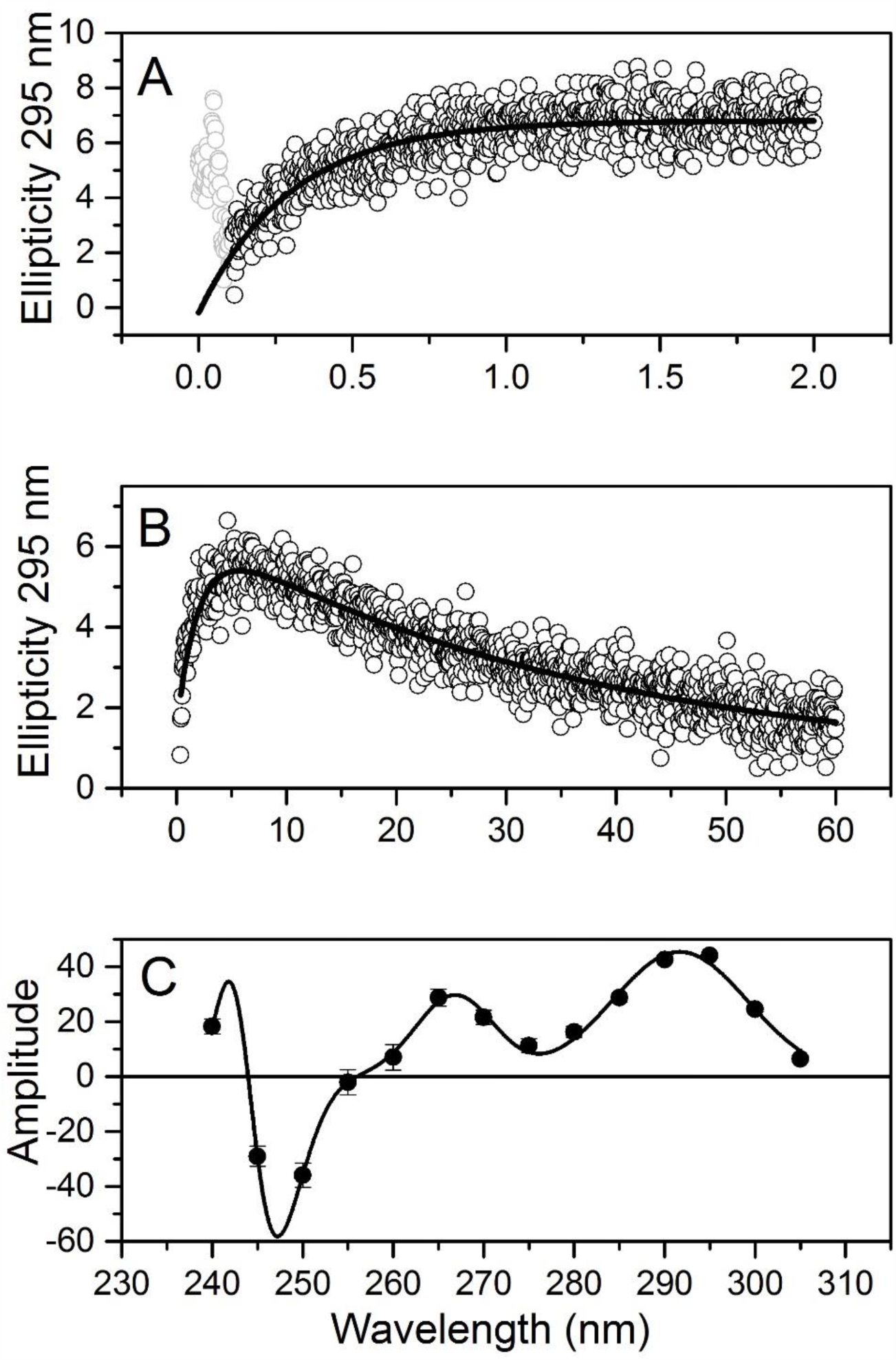
Single-wavelength stopped-flow studies of 1XAV folding. (A). CD change at 295 nm observed for 2 seconds after KCl mixing. (B) CD change at 295 nm observed for 60 seconds after KCl mixing. (C) Kinetic difference spectrum derived from stopped-flow measurements observed 2 seconds after KCl mixing.

An additional slower spectral change, indicative of a second intermediate form, was observed by hand mixing kinetic experiments with rapid wavelength scanning. Spectral scans were initiated within 10 s after KCl addition, and accumulated at 10 s intervals over 300 s. The results are shown in Figure 3A. These primary data show an initial transient spectral change near 295 nm that confirms the behavior seen in Figure 2A. Transient changes near 265 nm over a longer time scale are then observed. Singular value decomposition and nonlinear least-squares fitting of the amplitude vectors were used to analyze the data in Figure 3A. The best fit was obtained using a three-step sequential kinetic mechanism *U* ↔ *I*_1_ ↔ *I*_2_ ↔ *F*, where *U* and *F* are the unfolded initial and folded and final states, respectively, and *I_i_* are populated intermediate states. In addition to the relaxation times for the two steps seen in Figure 2, an additional slow step with a relaxation time ≥60 s was revealed by these experiments. The calculated spectra of the four species are shown in in Figure 3B. In the first step, *I*_1_ forms with a transient increase in CD at 295 nm, which then decreases as *I*_2_ forms, which has a maximum CD at 265 nm. Smaller CD changes at 265 nm then accompany the transition of *I*_2_ to the final folded form *F*. The time-resolved species plot is shown in Figure 3C. These kinetics are fully consistent with the single wavelength stopped-flow experiments shown in Figure 2. The overall folding reaction is complete by 200 s, with three resolved kinetic relaxation times of approximately 0.3, 9 and ≥60 s.

**Figure 3.**
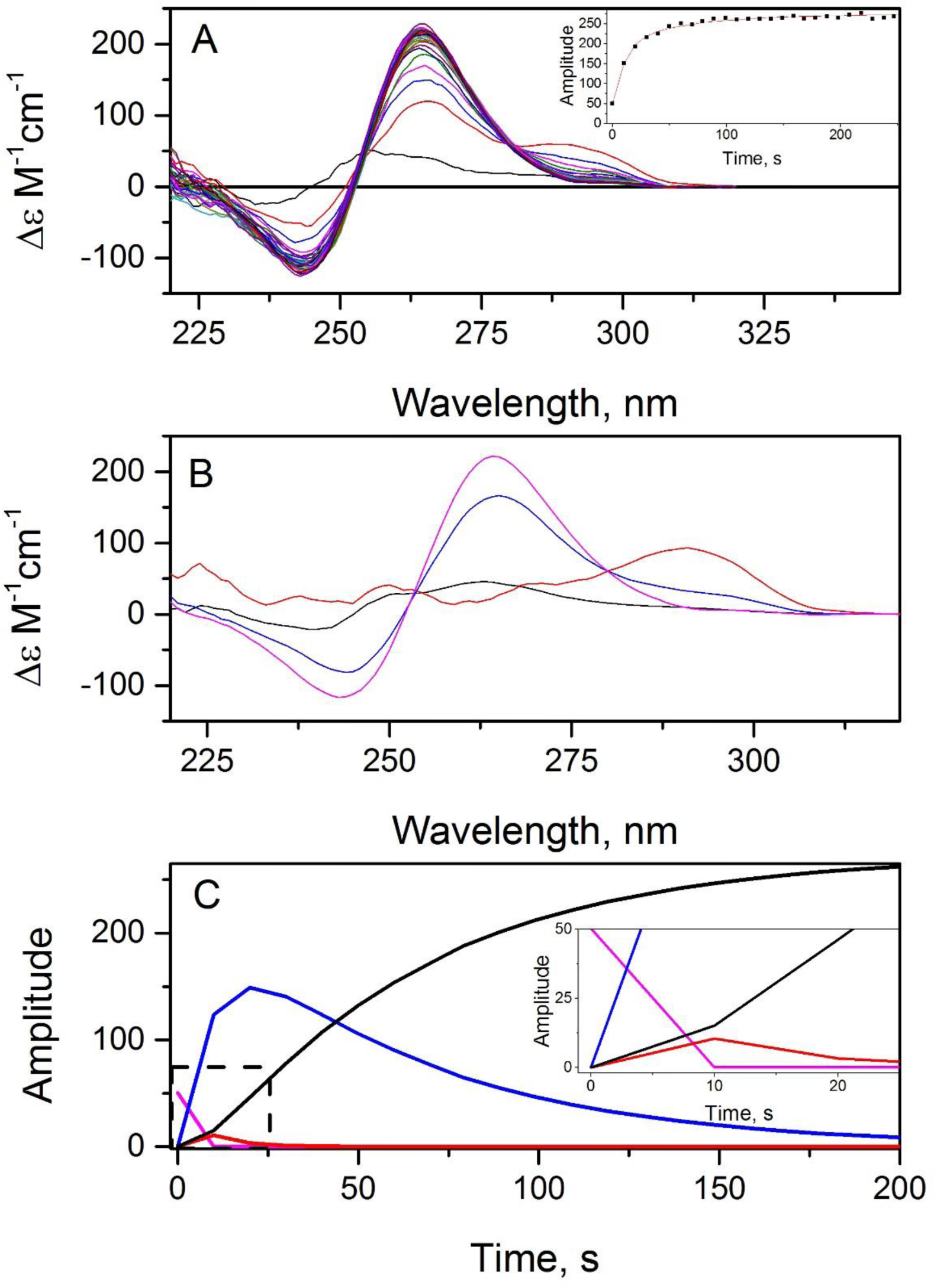
Kinetic studies of 1XAV folding using rapid scanning circular dichroism. (A) Primary data collected at 10 second intervals beginning 10 seconds after KCl mixing. The inset shows the time course of CD changes at 265 nm. (B) Computed CD spectra of kinetic species derived by SVD analysis. The black line is the initial unfolded strand. The red line is the first intermediate. The blue line is the second intermediate. The cyan line is the final folded 1XAV form. (C) Amplitudes of the changes in species spectra as a function of time as derived by SVD analysis. The inset shows an expanded scale for the initial portion of the time course. Colors refer to the spectral species shown in panel (B).

Rapid scanning stopped-flow absorbance studies (Figures S4-S5) and stopped-flow fluorescence studies using a 2-aminopurine substituted oligonucleotide (Figure S6) provided additional kinetic data that are fully consistent with the picture described above.

The rate of the first relaxation time was found to have a hyperbolic dependence on KCl concentration, while the second and third relaxation times were independent of KCl concentration up to 300 mM (data not shown). The temperature dependence of the first relaxation showed (anomalously) an apparent negative activation energy of approximately-11 kcal mol^−1^ (data not shown). Such behavior was previously observed for the folding of antiparallel G4 structures^1^ and indicates the presence of faster pre-equilibrium steps than can be observed by our methods.

We also characterized the kinetics of 1XAV unfolding by using the well-established complement trap method.^5, 20–21^ In the presence of complementary DNA, transiently unfolded G4 structures can be trapped as the more stable duplex, driving complete unfolding of the quadruplex. Since duplex formation is fast^33^, the observed slow apparent kinetics of duplex formation and G4 disappearance are assumed to arise from the rate-limiting unfolding of the G4 structure. We used ^1^H NMR to track unfolding kinetics using the procedure of Lane et al.^21^ Figure 4A shows 1D-^1^H NMR spectra of 1XAV obtained periodically during the course of complement-induced unfolding. The zero-time spectrum of folded 1XAV was identical to the 1-D proton NMR spectrum previously published by Ambrus et al.^26^ Comparison of our spectrum with that of Ambrus et al. allowed assignment of individual resonances to specific G residues. NMR spectra collected hourly over a period of 98 h after addition of 5-fold excess of 1XAV complementary DNA revealed slow disappearance of quadruplex-specific resonances with concomitant appearance of overlapping resonances in the duplex DNA region. The integrated intensities of quadruplex-specific resonances due to imino protons of G residues in the 10.5-12 ppm region are shown in Figure 4B. It is apparent that there was no preferential loss of signal due to specific G residues, (see Figure S7 for decay curves for individual resonances) implying that no specific region of the quadruplex unfolds separately on this time scale. The unfolding reaction required days to complete and followed a single exponential decay. An average time constant of 2.3 (± 0.3) × 10^5^ s (i.e. ≈ 63 h) was obtained by fits to the time-dependent changes in resonances for both individual G4 guanine residues and base pair formation. An independent experiment (Figure S7) using changes in fluorescence resonance energy transfer (FRET) for 5’-Fam, 3’-Tamra-labeled 1XAV, confirmed the slow unfolding kinetics seen by NMR.

**Figure 4.**
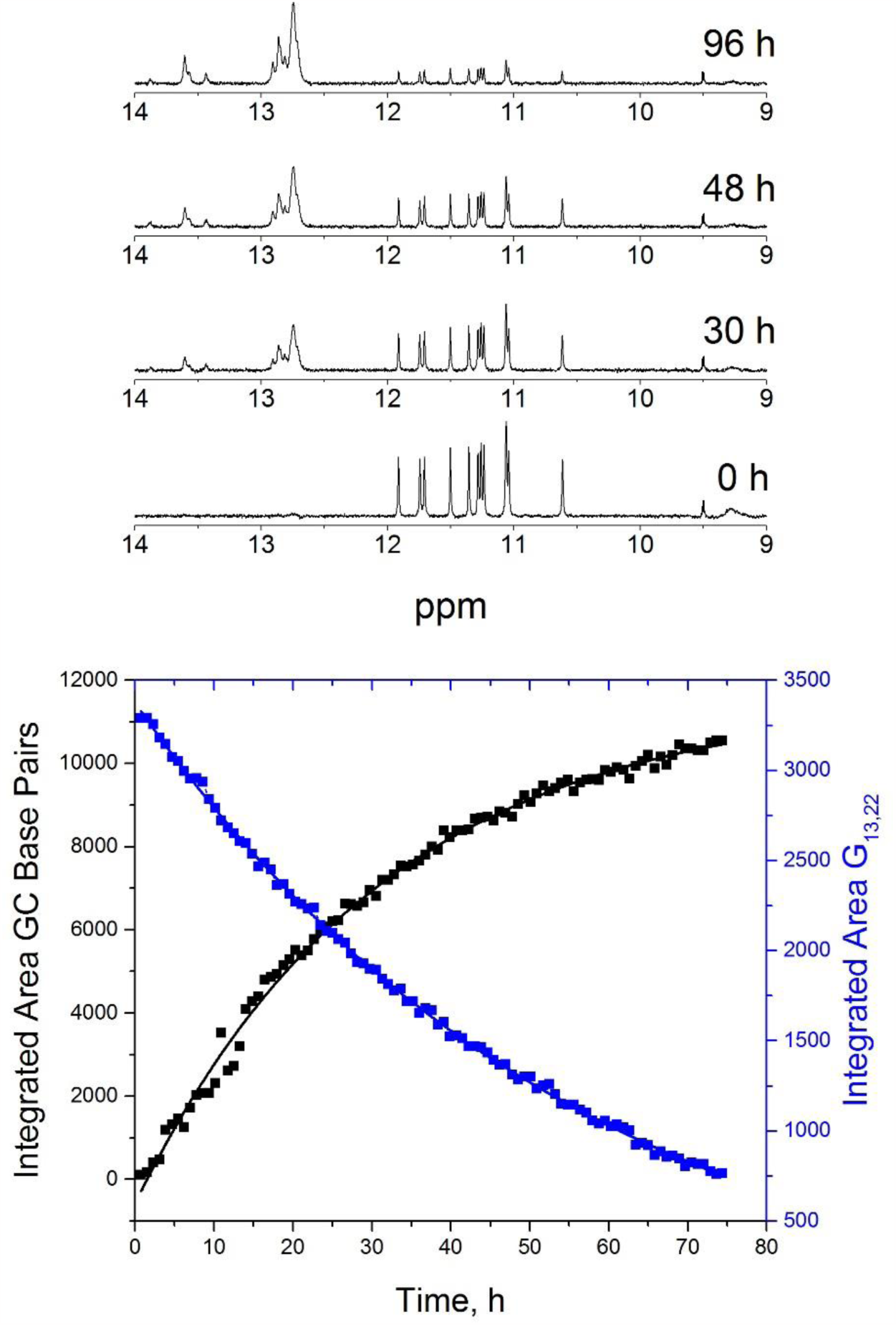
1XAV unfolding monitored by NMR. (Top) Selected NMR spectra obtained at the indicated times after addition of the complimentary trap strand. Resonances over the 10-12 ppm range are for guanines with the folded 1XAV structure. Resonances over the 12.5 – 14 ppm range are from duplex base pairs formed in the trap reaction.

Collectively, all of these kinetic experiments indicate the 1XAV folding is a complex process with populated intermediate steps. At a minimum, folding can be explained by a sequential three-step mechanism

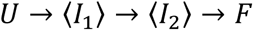

where the initial and final states are *U* and *F* and 〈*I*_i_〉 indicates populated ensembles of intermediates. Relaxation times of approximately 0.3, 2-9 and ≥60 s characterize the kinetic steps, and folding is complete within 200 s. Note that obligatory coupled K^+^ binding events are not specified in this simplified mechanism, but must be present. In addition, the KCl- and temperature-dependence of first relaxation time indicate that faster kinetic steps, beyond the resolution of our methods, surely exist. Only a single, very slow unfolding rate is observed. Because of the principle of microscopic reversibility, multiple additional unfolding steps must exist but are hidden by the slow rate-limiting process we observe.

A hypothetical model for G4 folding within the c-MYC promoter that is at least superficially consistent with the kinetic described here is shown in Scheme I. In the context of the duplex promoter, a DNA bubble must form (perhaps driven by supercoiling) to separate the strand containing the G4 forming sequence from its complementary strand. An antiparallel “chair” G4 structure might then form by way of a transient hairpin stabilized by G-G base pairing, a pathway similar that proposed for the early steps of the formation of telomeric G4 structures.^1-2, 5^ The spectral data in Figures 1 and 2 support the formation of such an intermediate. The final phase requires the unimolecular rearrangement of the antiparallel form to the final parallel structure.

To determine if there is a stereochemically feasible route from the chair to the final parallel form, a molecular dynamics simulation using the nudged elastic band method was used. This simulation involves setting the starting and final end points of the simulations and locating a low energy pathway between the two topological states. As the first experimentally observed intermediate has an antiparallel nature, a chair form of the 1XAV sequence d[G_3_TG_3_TAG_3_TG_3_] without 5ʹ and 3ʹ flanking bases was constructed and used as the starting structure. The corresponding truncated form of the known 1XAV parallel structure was the other endpoint. This exploration was not an extensive survey of the energy surface and low energy pathways, but since this method has not been previously applied to the chair to parallel topology transition, we wanted only to determine if the method could provide a possible pathway.

The transition from chair to parallel structure is a significant transformation because two diagonally opposed runs of guanines must change orientation and reverse direction in going from three lateral loops to three parallel loops (Scheme S1, Supporting Information). If these two transitions were simultaneous, the CD signature for G-quartets would disappear, which was not experimentally observed as the G-quartet CD and UV signature was maintained throughout. Therefore, the transition from chair to parallel topology could go via two pathways involving either parallel-parallel-lateral or lateral-lateral-parallel loop transitions (Scheme S1, Supporting Information). Both of these forms would have antiparallel or hybrid-like CD signatures with spectroscopic signals in the 295 nm region. These two forms were therefore created and inserted into the pathway in the nudged elastic band simulated annealing to guide the simulations. In both cases the simulations were able to successfully map a pathway from the chair to the parallel form of 1XAV, the results of which are visualized in movies provided in Supporting Information. A recent computational study^34^ simulated the folding pathway of a parallel RNA G4 structure and concluded, without any experimental verification, that the essence of the process is formation of compact coil-like structures stabilized by extensive cross-like interactions among the four G-strands. While such interactions may govern the fastest initial steps, our experimental results show that folded antiparallel structures are key intermediates along the folding pathway and must be taken into account.

The foregoing experiments may be relevant to the folding properties other promoter GQs. Bioinformatic analysis of the human genome suggests that transcriptional promotors are enriched in sequence motifs capable of folding into GQs.^35^ Based on characteristic CD spectra and DMS footprinting, many promoter sequences of oncogenes,^36^ including *c-MYC*, *VEGF*, *KRAS*, *c-kit*, as well as the promoter for human telomerase reverse transcriptase (*hTert*),^37^ contain substantial amounts of parallel GQ structures.

The c-*MYC* GQ consists of a 27-bp sequence (Pu27) with five G-tracts that lie upstream of the so-called P1 promoter which controls synthesis of the Myc protein, a transcription factor that is mutated in a variety of cancers.^3, 23, 38-40^ These mutations lead to constitutive expression of Myc and consequent dysregulation of cellular proliferation. It has been shown that GQ formation in Pu27 prevents *c-MYC* expression; thus characterization of factors influencing the stability of Pu27 model sequences may be relevant to strategies for controlling aberrant Myc activity. The timescale of the folding of 1XAV we observe is relevant since the cell cycle of a mammalian cell is roughly 1 day, and the transcription of a typical gene is roughly 10 minutes.^41^ In contrast, the extremely slow unfolding rate that we observe seems to be too slow to be of any physiological relevance, suggesting that disruption of G4 structures in vivo may need catalytic assistance. Indeed, such enzymes exist. Numerous G4 unwinding helicases have been identified that target and regulate specific G-quadruplex structures^42–43^ that would serve to accelerate the slow unfolding reaction we observe.

## ASSOCCIATED CONTENT

### Supporting Information

The following files are available free of charge. A PDF file containing Experimental Methods, two Tables, seven Figures and one Scheme. Two mp4 files showing movies of the transition from the chair to a parallel G-quadruplex conformations derived from nudged elastic band simulations.

### Notes

The authors declare no competing financial interests.

## ACKNOWLEDGMENT

Supported by grants CA35635 and GM077422 from the National Institutes of Health.

---

**Scheme 1.**
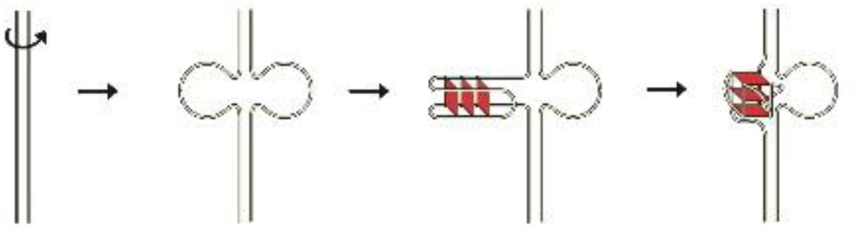
Hypothetical folding pathway within the c-myc promoter, as described in the text.

