## Supporting Information for "Folding landscape of a parallel G-quadruplex"

**J. G. Brown Cancer Center, University of Louisville, 505 S. Hancock St., Louisville, KY 40202**

### Supporting Information

#### Materials

The oligodeoxynucleotide sequences and their extinction coefficients used in this study are listed in Table S1. Unlabeled oligos were obtained from Integrated DNA Technologies, Coralville, IA as lyophilized gel-purified products. 5'-FAM, 3'-Tamra (FRET)-labeled 1XAV was from Eurofin Genomics, Louisville, KY and was HPLC-purified. Oligos were reconstituted in deionized water at concentrations of 1-3 mM and stored at 4 °C. Oligonucleotide concentrations were estimated from their absorbance at 260 nm determined on suitable dilutions into 10 mM $LiPO_4$, 1 mM EDTA, pH 7.0 (Li-buffer). To prepare working solutions, the stock was diluted into Li-buffer, (with or without supplemental KCl), heated for 10-20 min in a 1-L boiling water bath followed by slow overnight cooling to room temperature. Solutions for NMR were dialyzed overnight vs. Li-buffer with 25 mM KCl but without EDTA. Other reagents were from Sigma (KCl), Aldrich ($LiH_2PO_4$ and LiOH) and Fluka (EDTA acid).

#### Methods

*CD Spectroscopy.* UV circular dichroism and absorption spectra were measured at 25 °C in 1-cm cuvettes with a Jasco J-810 spectropolarimeter equipped with a Peltier temperature-controlled sample holder. Unless noted otherwise, spectra were scanned from 340 to 220 nm at a rate of 200 nm/s with a 1 nm spectral bandwidth and a digital integration time of 2 s. Four replicate spectra were averaged. Blank-corrected spectra were normalized using the relationship $\Delta\varepsilon = \theta/(32980 \cdot c \cdot l)$ where $\theta$ is the observed ellipticity in millidegrees, $c$ is the molar strand concentration and $l$ is the pathlength in cm.

*Equilibrium Titrations with KCl.* Prior to initiating the kinetic studies, titrations of unfolded 1XAV with KCl were carried out to establish the spectroscopic properties of the unfolded and equilibrium folded states of 1XAV in the presence of this cation. Initially, these titrations were conducted in tetrabutyl ammonium phosphate buffer (tBAP), a buffer system that did not appreciably promote folding of telomeric oligonucleotides such as Tel22 (d[A(GGGTTA)$_3$GGG]) in the absence of $K^+$ or $Na^+$.[1-2] In contrast to these telomeric sequences, 10 mM tBAP appeared to promote folding of 1XAV in the absence of $K^+$ (based on CD data not shown). To avoid this complication, kinetic and equilibrium experiments were carried out in buffer consisting of 10 mM $LiPO_4$, 1 mM EDTA, pH 7.0, which, based on CD criteria, did not promote quadruplex formation.

KCl binding isotherms for 1XAV were constructed by mixing an aliquot of KCl from a concentrated stock solution with heat-denatured, annealed 1XAV (5.5 µM) in a 1-cm cuvette. After mixing, the cuvette was placed in the sample compartment of the spectropolarimeter and equilibrated for 3 min at 25 °C, after which four replicate spectra were recorded. Each point on the titration curve was obtained with a unique solution. CD spectra of these solutions measured after overnight incubation were identical to those obtained soon after preparation. The baseline-corrected, normalized data matrices consisting of CD spectra vs. [KCl] were analyzed by singular value decomposition (SVD) using routines in Matlab 7.1 (MathWorks, Natick, MA). The titration curve shown Figure 1A was fit to the biphasic dose-response curve equation available in GraphPad Prism version 6.07 for Windows (GraphPad Software, Inc., LaJolla, CA, USA).

*Kinetic experiments*. The kinetics of $K^+$-induced folding of 1XAV was assessed by stopped-flow and manual mixing methods. Rapid changes in absorbance were monitored as previously described[1] using an Olis (On-Line Instrument Systems, Bogart, GA) rapid scanning stopped-flow spectrophotometer equipped with a water bath for temperature regulation. The Olis-supplied software (GlobalWorks) produces a data matrix **Y** consisting time-resolved difference spectra $\Delta A_{i,j}$ where row *i* represents $\lambda_i$, column *j* represents time $t_j$, and $\Delta A = A_{Unfolded} - A_{Folded}$. Analysis of **Y** by SVD reduces it to the product of 3 matrices: **U** which contains the kinetic information, **V** which contains the spectral information, and the diagonal matrix **S** which contains the singular values $S_{i,j}$ (the relative weight of each component). The significant kinetic eigenvectors representing the time-dependent changes in absorption spectra between ~260 and ~320 nm were fit by non-linear least squares to the expression $\Delta A_t = \Sigma exp(-k_i*t)$ where *i* is the number of significant components derived from the SVD analysis of the data set. The fitting process generates the absolute spectra of the kinetically significant species as well as their evolution over time.

Rapid changes in CD were determined using the stopped-flow CD attachment for the Jasco J810 as previously described[2]**.** The pathlength of the observation cell was 0.2 cm and reactions were observed at room temperature (~22.5 °C). The progress of folding was monitored in individual kinetic experiments by recording changes in CD signal at 5-nm intervals over the wavelength range 240-305 nm. Depending on the noise level, from 5 to 10 successive progress curves were averaged at each wavelength. The amplitudes and time constants describing the kinetic data at each λ were then determined by fitting the progress curves to one or two exponentials using the non-linear least squares routines in Origin 7.1. For some of the noisier data, the relaxation time for the reaction was held at a constant value determined from less noisy data and only the amplitude term was adjusted.

*Unfolding Kinetics*. Unfolding kinetics of 1XAV was assessed by the complement trapping method using 1D-proton NMR to measure the appearance of duplex and disappearance of GQ resonances. The experiments were carried out at 25 °C essentially as described by Lane et al.[3] This method takes advantage of the separation of Hoogsteen-bonded guanine imino proton resonances in the quadruplex state located at 12-10.5 ppm from the AT and GC resonances in the duplex found at 14-12.5 ppm. The unfolding reaction was initiated by adding a 5-fold molar excess of 1XAV complementary DNA to 0.3 mM 1XAV prefolded in 25 mM KCl. 1D proton NMR spectra were acquired on an 800 MHz Varian Unity Inova NMR spectrometer using 3-9-19 watergate pulse sequence to suppress the dominant water peak with a delay of 100 µs between the 3-9-19 pulses. A spectral width of 20000 Hz, an acquisition time of 1.5 s and a relaxation delay of 1.5 s were used for a total of 256 scans for signal averaging. A pre-acquisition delay of about 2816 s was introduced after the first measurement so that each measurement was carried out approximately every hour for 96 hours. The decrease in the peak area of the quadruplex resonances and the increase in peak area of the duplex resonances were quantified using the program Mnova 9 (Mestrilab Research, Santiago de Compostela, Spain).

*Molecular dynamics*. The chair to parallel transition was modeled off the 1XAV NMR structure[4] by the nudged elastic band method as implemented in the AMBER 16 program Sander. The flanking bases were removed and the sequence used was d[$G_3TG_3TAG_3TG_3$]. The chair form was manually created by adjusting the polarity of the second and fourth G-run flipping the bases to maintain the three G-tetrads. Inter-tetrad coordinated potassium ions were retained. The corresponding parallel form was the 1XAV structure without flanking bases. The initial endpoints were explicitly solvated and potassium counter ions were generated with the parm16SB.dat Amber force field using the following protocol: (i) two unsolvated potassium ions were placed between the G-quartet tetrads for stabilization (ii) the system was solvated by the addition of a rectangular box of TIP3P water at 15 angstroms, and (iii) neutralizing solvated potassium ions were added randomly around the quadruplex structures using Amber 12 leap rules for counter ions. Energetically stable models were generated using the following protocol: minimize water holding the DNA (50 kcal $mol^{-1}$ $Å^{-1}$), minimize the complete system, 50 ps molecular dynamics (heating to 300 K) holding the DNA fixed (50 kcal $mol^{-1}$ $Å^{-1}$), (iv) unrestrained molecular dynamics for 10 ns. Simulations were performed in the isothermal isobaric ensemble (P = 1 atm, T = 300 K). Periodic boundary conditions and the Particle-Mesh-Ewald algorithm were used. A 2.0 fs time step was used with bonds involving hydrogen atoms frozen using SHAKE. For the equilibration steps and the production steps, molecular dynamics calculations were carried out using AMBER 16 program sander and the cuda version of pmemd, respectively. The final preparation step was minimization with implicit solvent, igb=1, saltcon=0.2.

The nudged elastic band method using simulated annealing was used to map a pathway from chair to the parallel form using multisander as per the AMBER website and using our published protocol (2). 256 images were used and tgtfitmask=":1-16" including the tetrad core potassium ions and tgtrmsmask=":1-16@P,O1P,O2P,O5',O3'" were used. The endpoints were set as the chair and corresponding hybrid form. The five steps were i) heating: linear heating from 0 to 300K for 20ps with a 0.5fs time step with neb options skmin = 10,skmax = 10, ii) equilibrium: 300K 100ps with a 1fs timestep with skmin = 50,skmax = 50 used and in subsequent steps, iii) simulated annealing: 600ps, 1fs time step, 0-50ps heat 300K-400K, 50-100ps 400K, 100-150ps heat 400K-500K, 150-200ps 500K, 200-250ps cool to 300K, 250-300ps 300K, iv) slow cooling: 120ps with 1fs time step 300K-0K, v) long cool: 200ps 0K with skmin = 10,skmax = 10. The final pathway was extracted and visualized using a modified combine final pathway.sh script (Supplementary movie1 and 2).

**Supporting Tables**

Table S1. Oligodeoxynucleotides used in this study

| Name | Sequence | ε ($mM^{-1}$ $cm^{-1}$) |
|---|---|---|
| 1XAV | 5'-TGA GGG TGG GTA GGG TGG GTA A-3' | 228.7 |
| C-1XAV | 5'-TTA CCC ACC CTA CCC ACC CTC A-3' | 191.5 |
| 1XAV-12AP | 5'-TGA GGG TGG GT2-aminopurine GGG TGG GTA A-3' | 215.2 |
| Fret-1XAV | 5'-Fam-TGA GGG TGG GTA GGG TGG GTA A-3'-Tamra | 277.6 |

Fam = 6-carboxyfluorescein; Tamra = 5-Carboxytetramethylrhodamine

Table S2. Fitted parameters for kinetic data in Figure S3.[a]

| Wavelength (nm) | y0 (mdeg) | Std Error | Amplitude (mdeg) | Std Error |
|---|---|---|---|---|
| 305 | -0.14 | 0.03 | -1.02 | 0.14 |
| 300 | 3.14 | 0.03 | -3.90 | 0.16 |
| 295 | 6.81 | 0.03 | -6.99 | 0.14 |
| 290 | 8.55 | 0.03 | -6.74 | 0.16 |
| 285 | 9.71 | 0.05 | -4.56 | 0.24 |
| 280 | 11.25 | 0.06 | -2.58 | 0.32 |
| 275 | 14.68 | 0.08 | -1.80 | 0.40 |
| 270 | 19.52 | 0.08 | -3.42 | 0.41 |
| 265 | 22.16 | 0.09 | -4.55 | 0.49 |
| 260 | 21.89 | 0.14 | -1.12 | 0.74 |
| 255 | 10.26 | 0.14 | -0.32 | 0.72 |
| 250 | 0.12 | 0.14 | 5.68 | 0.70 |
| 245 | -3.96 | 0.11 | 4.59 | 0.57 |
| 240 | -2.66 | 0.09 | 2.88 | 0.44 |

[a]Data sets in Figure S3 were fit to a single exponential with a fixed relaxation time τ of 300 ms.

**Supporting Figures**

**S1. Equilibrium titration of 1XAV with KCl is a three-step reaction.**

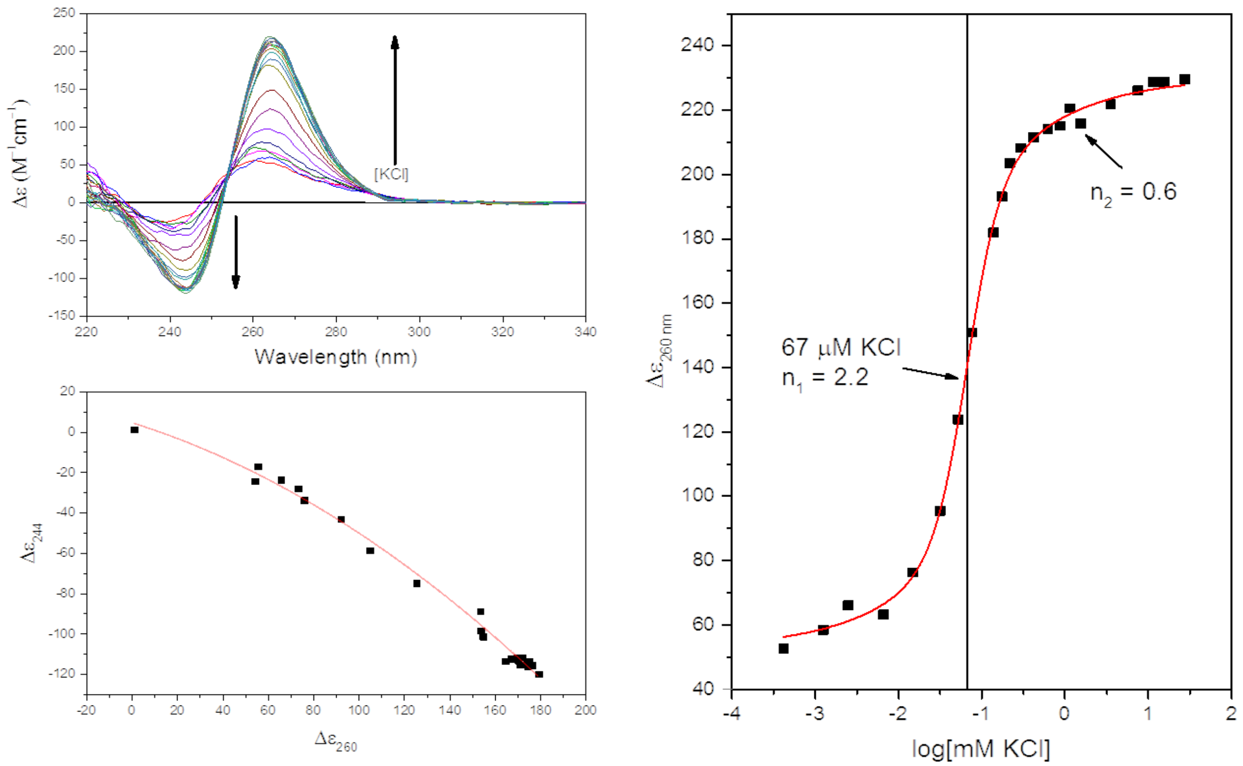


**Figure S1.** *Top Left*: titration of 1XAV with KCl assessed by CD. Conditions: [1XAV] = 5.5 µM in 10 mM $LiPO_4$ buffer, 1 mM EDTA, pH 7.0, 25 °C. *Bottom Left*: plot of the CD signal at 244 nm vs. the signal at 260 nm. As explained in the text, the non-linear relationship indicates that more than two spectroscopic species are present during the titration. *Right*: non-linear least squares fit of the molar ellipticity at 260 nm to a double Hill plot.

S2. **Thermal denaturation of 1XAV is a three-step process.**

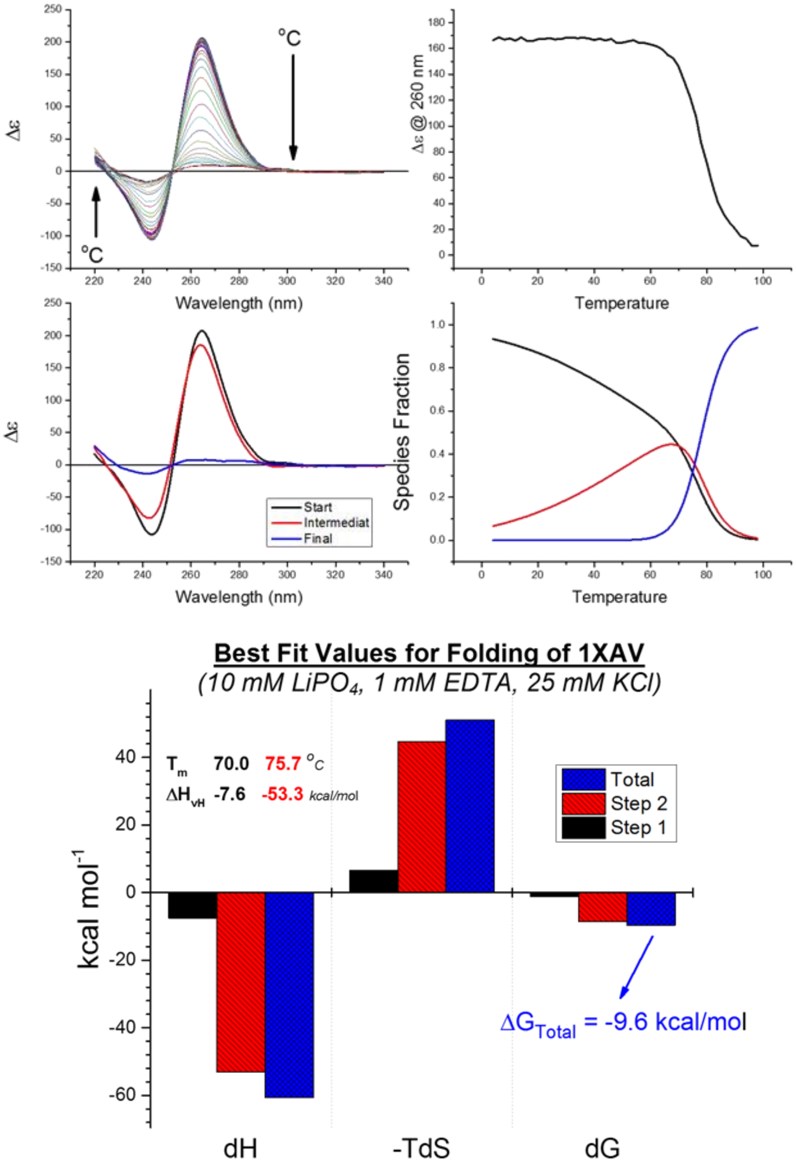


**Figure S2.** SVD analysis of temperature-dependent CD spectra of 1XAV in 10 mM $LiPO_4$ buffer, 1 mM EDTA, 25 mM KCl. *Top Left*: normalized CD curves measured as a function of sample temperature. *Top Right:* Dependence of Δε(260 nm) on temperature. SVD analysis (not shown) of the temperature-wavelength data matrix indicated that the data set was comprised of three spectrally significant species. Fitting to a single intermediate, 2-step mechanism F ↔ I ↔ U allowed calculation of the CD spectra of the three species (*Middle Left*), the midpoint temperature $T_m$ for each, the enthalpy change associated with each step, and the concentration profiles of each species as a function of temperature (*Middle Right*). The *Bottom* panel depicts a complete thermodynamic analysis of the thermal stability of 1XAV in 25 mM KCl.

**S3. Single wavelength stopped-flow kinetic studies reveal the kinetic CD spectrum during the initial 2s of the folding reaction.**

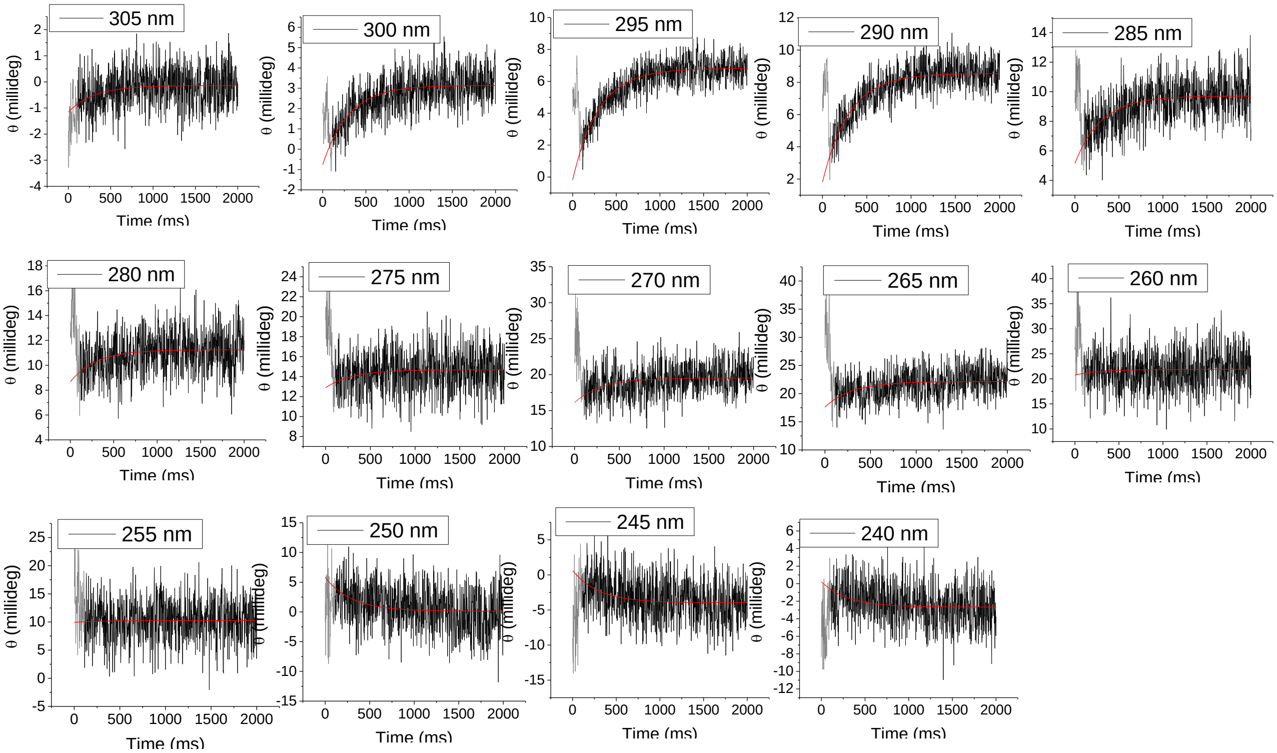


**Figure S3.** Wavelength dependence of the kinetics of KCl-induced folding of 1XAV determined by stopped-flow CD. To determine the amplitude of the CD change, the CD signal was fit to a single exponential with a fixed relaxation time constant τ = 300 ms. The θ values shown in grey represent the CD signal recorded during flow and were not included in the least squares fit. Conditions: 24 µM 1XAV, 25 mM KCl, 10 mM $LiPO_4$, 1 mM EDTA, pH 7.0, room temperature (~23 °C). Reagent concentrations are after 1:1 mixing. Observation pathlength = 0.2 cm. The least squares optimized fitting parameters for each data set are summarized in Table S2.

**S4. Rapid scanning stopped-flow absorbance experiments suggest a complex folding process for 1XAV and reveal an antiparallel intermediate on the pathway to formation of an all-parallel final structure.**

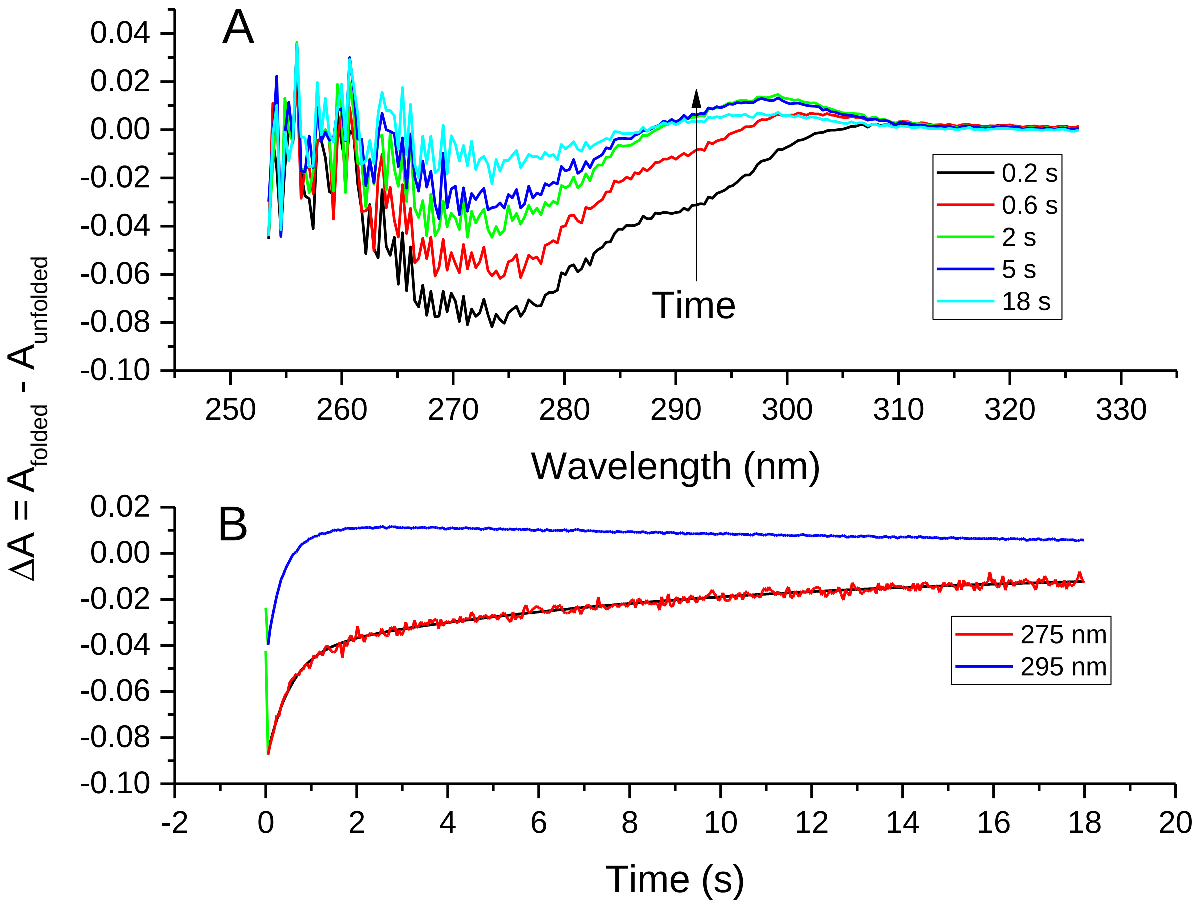


**Figure S4**. Kinetics of 1XAV folding observed by wavelength scanning stopped-flow. Panel A shows difference spectra determined at the indicated times after 1:1 mixing 7.4 μM 1XAV with 50 mM KCl at 25 °C. Absorbance changes (ΔA) were measured with reference to the absorbance of folded 1XAV determined several minutes after mixing. The lower panel B compares time-dependent changes in absorbance at 275 nm and 295 nm. The different kinetic profiles at these wavelengths emphasizes the presence of kinetic intermediates with different UV spectra.

**S5. Analysis of kinetic data shown in Figure S4 by singular value decomposition and nonlinear fitting.**

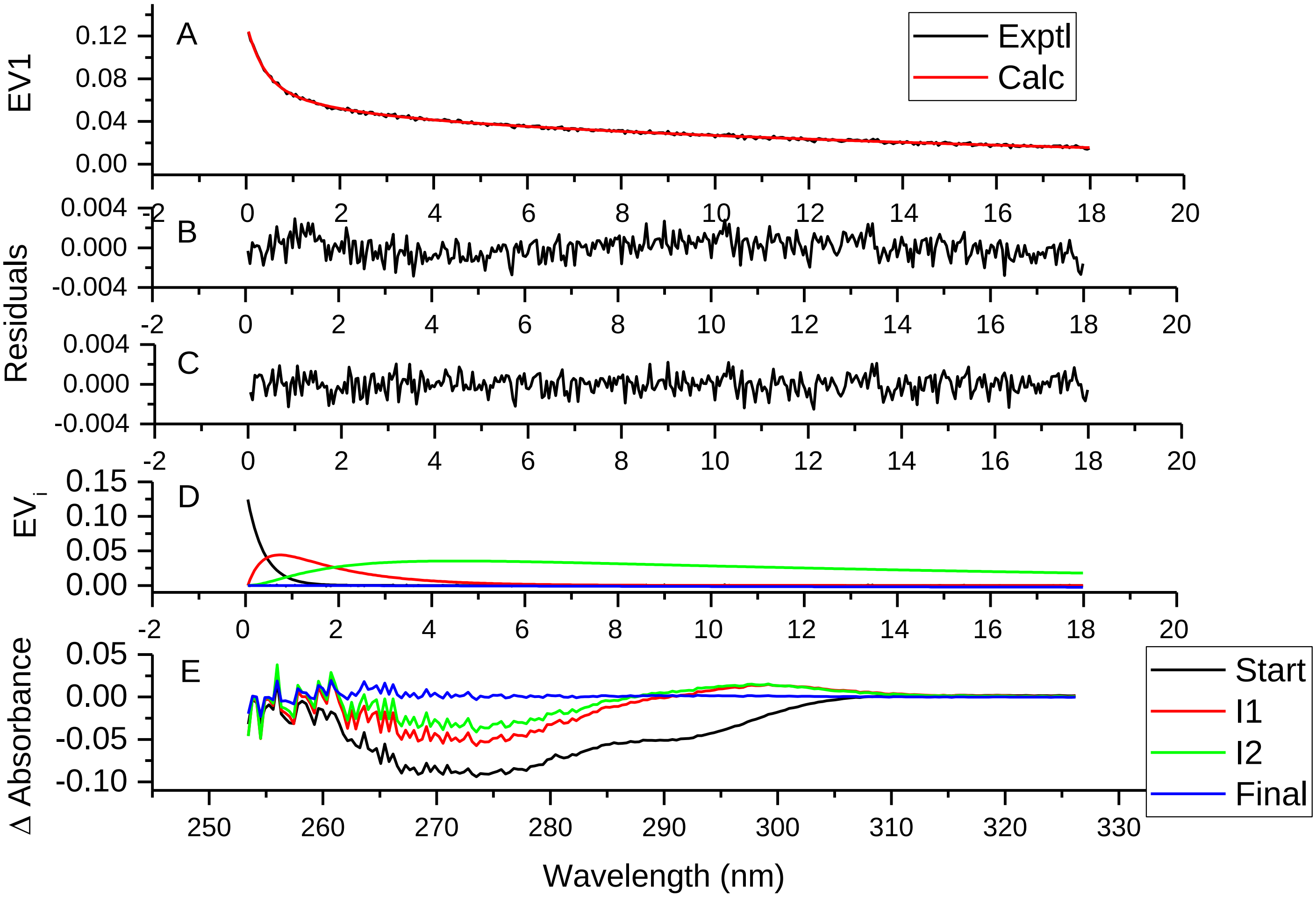


**Figure S5**. SVD analysis of kinetic difference spectra in Figure 1 shows that four spectrally distinct species contribute to ΔA at various times. The best fit of the data set was obtained with a reaction pathway with two intermediates: U ↔ I1 ↔ I2 ↔ F. Panel A shows a least squares fit of the most significant eigenvector to the three-exponential expression describing the two-intermediate mechanism. The black points represent the experimental data, the red line is the best fit. Panels B and C show the residual plots for 1- and 2-intermediate fits. Panel D shows the calculated eigenvector absorbance profiles of the U, I1, I2 and F species. Panel E shows the calculated difference spectra of the species U, I1, I2 and F.

**S6. Stopped-flow fluorescence with 1XAV-12AP is a two-step process**.

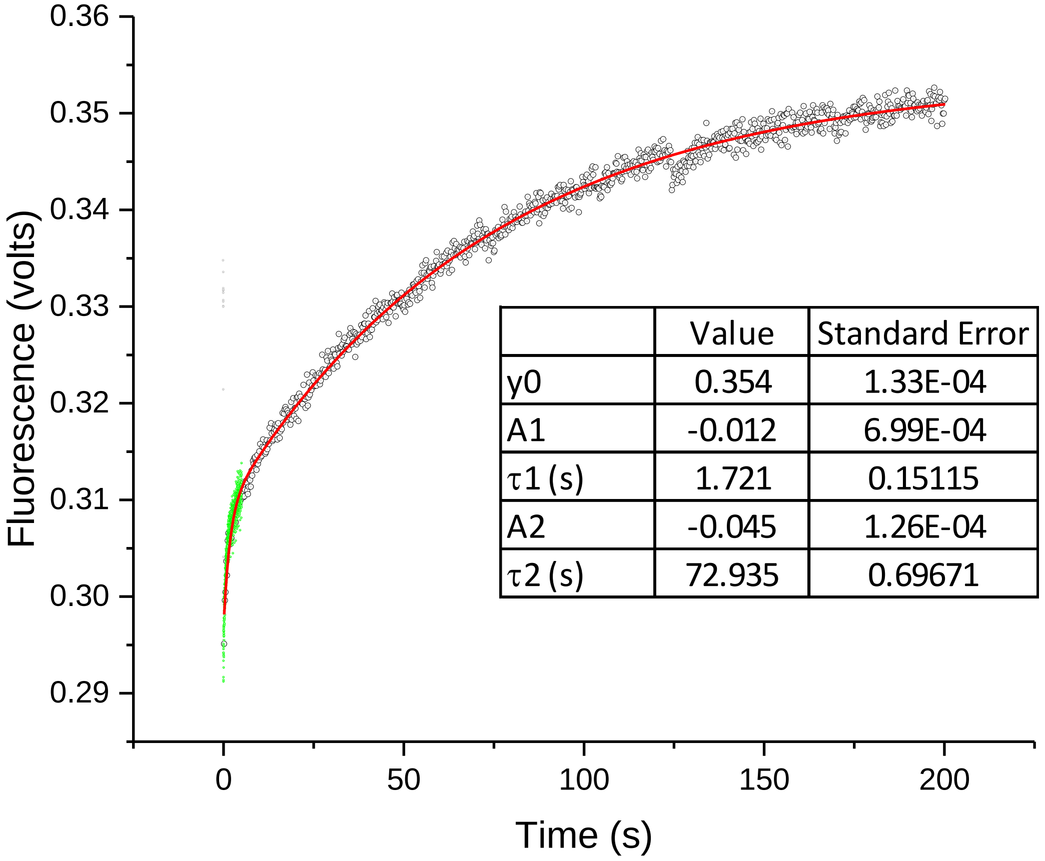


**Figure S6.** Folding kinetics of 1XAV-12AP. 2-Aminopurine fluorescence was excited at 305 nm and fluorescence emission was detected at 90° through a 320-nm cufoff filter using the OLIS stopped-flow apparatus in fluorescence mode. Conditions: [1XAV-12AP] = 1.5 µM, [KCl] = 25 mM in 10 mM $LiPO_4$ buffer, 1 mM EDTA, pH 7.0, 25 °C. The black and green points show experimental data collected independently at a rapid and slow rate. The grey points are data collected during mixing and flow and were omitted from the fit. The red line represents the best fit of the data to a sum of two exponentials.

**S7. Unfolding of FRET-labelled 1XAV in the presence of 1XAV-complement is slow, confirming NMR results.**

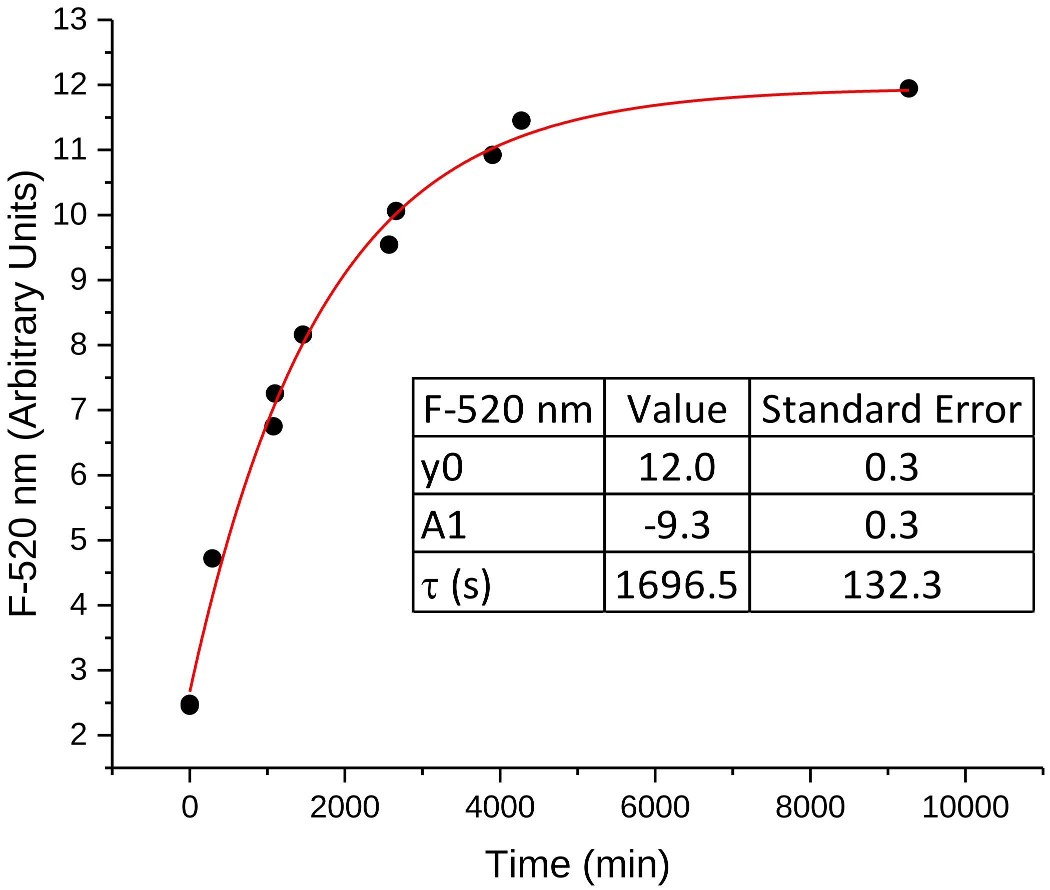


**Figure S7.** Unfolding of 5'-Fam-1XAV-3'Tamra (FRET-1XAV, 0.5 μM) in the presence of a 5-fold molar excess of 1XAV-complement. Conditions: 10 mM $LiPO_4$, 1 mM EDTA, 25 mM KCl, pH 7.0, 25C. The points represent the experimental data and the red line shows the best fit of the data to a single exponential using the optimized parameters in the inset.

**S8. Model for the chair-to-parallel transition.**

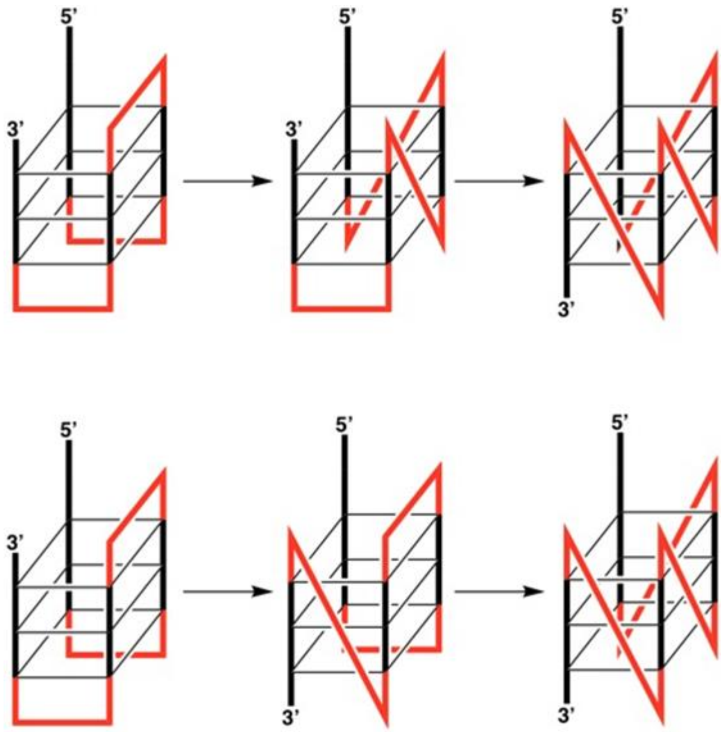


Scheme SI. Two possible pathways for the chair to parallel G4 conformation transition. Loops are show in red. (Top) The transition begins near the 5’ end with the second G-run inversion of direction and (Bottom) the transition begins with the 3’ G-run inversion of direction.
